# A biodiverse package of southwest Asian grain crops facilitated high-elevation agriculture in the central Tien Shan during the mid-third millennium BCE

**DOI:** 10.1101/2020.02.06.936765

**Authors:** Giedre Motuzaite Matuzeviciute, Taylor R. Hermes, Basira Mir-Makhamad, Kubatbek Tabaldiev

**Affiliations:** Department of Archaeology, Vilnius University, Vilnius, Lithuania

## Abstract

We report the earliest and the most abundant archaeobotanical assemblage of southwest Asian grain crops from Early Bronze Age Central Asia, recovered from the Chap II site in Kyrgyzstan. The archaeobotanical remains consist of thousands of cultivated grains dating to the mid-late 3^rd^ millennium BCE. The recovery of cereal chaff, which is rare in archaeobotanical samples from Central Asia, allows for the identification of some crops to species and indicates local cultivation at 2000 m.a.s.l., as crop first spread to the mountains of Central Asia. The site’s inhabitants cultivated two types of free-threshing wheats, glume wheats, and hulled and naked barleys. Highly compact morphotypes of wheat and barley grains represent a special variety of cereals adopted to highland environments. Moreover, glume wheats recovered at Chap II represent their most eastern distribution in Central Asia so far identified. Based on the presence of weed species, we argue that the past environment of Chap II was characterized by an open mountain landscape, where animal grazing likely took place, which may have been further modified by people irrigating agricultural fields. This research suggests that early farmers in the mountains of Central Asia cultivated a high diversity of southwest Asian crops during the initial eastward dispersal of agricultural technologies, which likely played a critical role in shaping montane adaptations and dynamic interaction networks between farming societies across highland and lowland cultivation zones.

## Introduction

Crops domesticated in various locations of Eurasia spread widely to new environments unlike those where they were initially cultivated. A number of crops originating in diverse landscapes of what is now present-day China, such as millets, hemp and buckwheat, spread to Europe, while a variety of southwestern Asian crops, such as wheat and barley, became important food sources across monsoonal Asia [1–4]. In the past decade, sampling for botanical remains during archaeological excavation has become routine, and subsequent analysis has transformed our understanding of the timing and routes of plant dispersals through Central Asia [5,6]. Along these lines, one of the most important discoveries at the forefront of this fluorescence in archaeobotanical research in Central Asia was the earliest millet and wheat found together at the site of Begash in the Dzhungar Mountains of Kazakhstan dated to the end of the 3^rd^ millennium BCE [5,6]. Further work across China showed that during the first half of the second millennium BCE, wheat and barley became fully integrated into millet cultivation systems, thus transforming agricultural strategies and culinary traditions of communities in ancient China [4,5,8–10].

Many questions still remain, however, in understanding the status of domesticated crops in diverse societies across Eurasia during the earliest stages of multidirectional agricultural dispersals [1,11–13]. We especially have a poor understanding of what species and their morphotypes were moved and why [14,15], and whether different crop species took the same pathways across wide swaths of Asia [4]. An additional enigma is represented by the earliest broomcorn millet and bread wheat from Begash that were found exclusively in a human cremation cist without crop processing chaff [7]. As this mortuary context does not inform whether these grains were consumed by people or locally cultivated, Frachetti [12] applied the term “seeds for the soul” to describe a cultural context of the seeds that was focused on ritual use. Recently, isotopic analysis employing incremental sampling of ovicaprid tooth enamel bioapatite for δ^13^C and δ^18^O values has shown that domesticated animals at Begash and the adjacent Dali site, the latter dating approximately 400 years earlier than the former, were foddered with millet during winter, suggesting that the spread of pastoralist subsistence eastward was intricately tied to the subsequent westward dispersal of millet [16]. While this research sheds light on the intersection of the ritual use and economic value of millets, the cultural status of southwest Asian crops for local Central Asian societies during this period of early crop dispersals remains unclear.

Whether wheat and barley constituted a singular crop package during their dispersal remains an important line of inquiry. Current evidence suggests that barley took a different pathway than wheat to reach the present territory of China, as the radiocarbon dates measured on wheat and barley are substantially different across China, revealing that barley had spread later by people along the highland route of the Inner Asian Mountain Corridor and the southern regions of the Himalayan Mountains [4]. It has also been noted that small morphotypes of wheat were selected for cultivation as wheat moved eastward to China, since diminutive and rounder wheats may have been better suited to the culinary traditions dominated by small-grained Chinese millets [14]. On the contrary, additional research has argued that compact morphotypes of both wheat and barley recovered archaeologically in environmental margins, such as the highlands of Central Asia is strongly influenced by environmental adaptation rather than culinary preferences [15,17–19].

It is important to emphasize that our current understanding about the initial stages of the Eurasian crop interchange across Central Asia is based on an extremely small quantity of plant remains recovered from only a few archaeological sites dated to the 3^rd^ millennium BCE, such as Begash and Tasbas in Kazakhstan, Sarazm in Tajikistan, and South Anau in Turkmenistan [20–23]. For example, at Begash (2450–2100 cal BCE), cultivated crops are represented only by 13 compact free-threshing wheat grains, one barley grain, and 61 grains of broomcorn millet, while at Tasbas (2840–2496 cal BCE) just five compact free-threshing wheat grains were found, in addition to 11 ambiguously identified “Cerealia” grains [18,23]. Such a small quantity of domesticated plant grains per site does not allow us to understand cultivation strategies, the relative importance of consumption versus ritual use, or the coherence of early agricultural packages.

Here, we present an early archaeobotanical assemblage consisting of five cultivated crops of southwest Asian origin recovered from the high-elevation Chap II site located in the central Tien Shan Mountains of Kyrgyzstan. The archaeobotanical samples represent the largest crop assemblage so far found at 3^rd^ millennium BCE sites between the Pamir, Tien Shan, and Altai Mountains, representing thousands of carbonized cereal remains. The assemblage reflects a strikingly high degree of biodiversity in wheat and barley grain morphotypes. Analysis of the accompanying weed taxa recovered from Chap II hints on the past ecology of cereal cultivation, suggesting that crops may have been grown with the aid of irrigation, while recovered chaff fragments indicate local cultivation and processing. Notably, we present chaff and grain remains of glume wheat that, for the first time, were recovered from the eastern regions of Central Asia.

## The Chap II site

The Chap II site is located in the Kochkor valley of central Kyrgyzstan at 2000 m.a.s.l. (42°10’51.7” N, 75°51’3.64” E). The site is situated inside the inundation of a small loess hill that protrudes from the foothills into the southeastern part of the 80-km long Kochkor valley (Fig 1). The archaeological remains of Chap II were recovered through excavation of the Chap I farmstead (1065-825 cal BCE), which was occupied during the transitional period from the Late Bronze Age to the Early Iron Age [24]. In previous years, a small 6.5×6.5 m trench was excavated that yielded a diverse assemblage of ceramic sherds, crop processing tools, domestic animal skeletal remains, carbonized remains of domesticated grains and chaff, which together reflect extensive agricultural and pastoralist activities by the inhabitants of the Chap I site [24].

**Fig 1.**
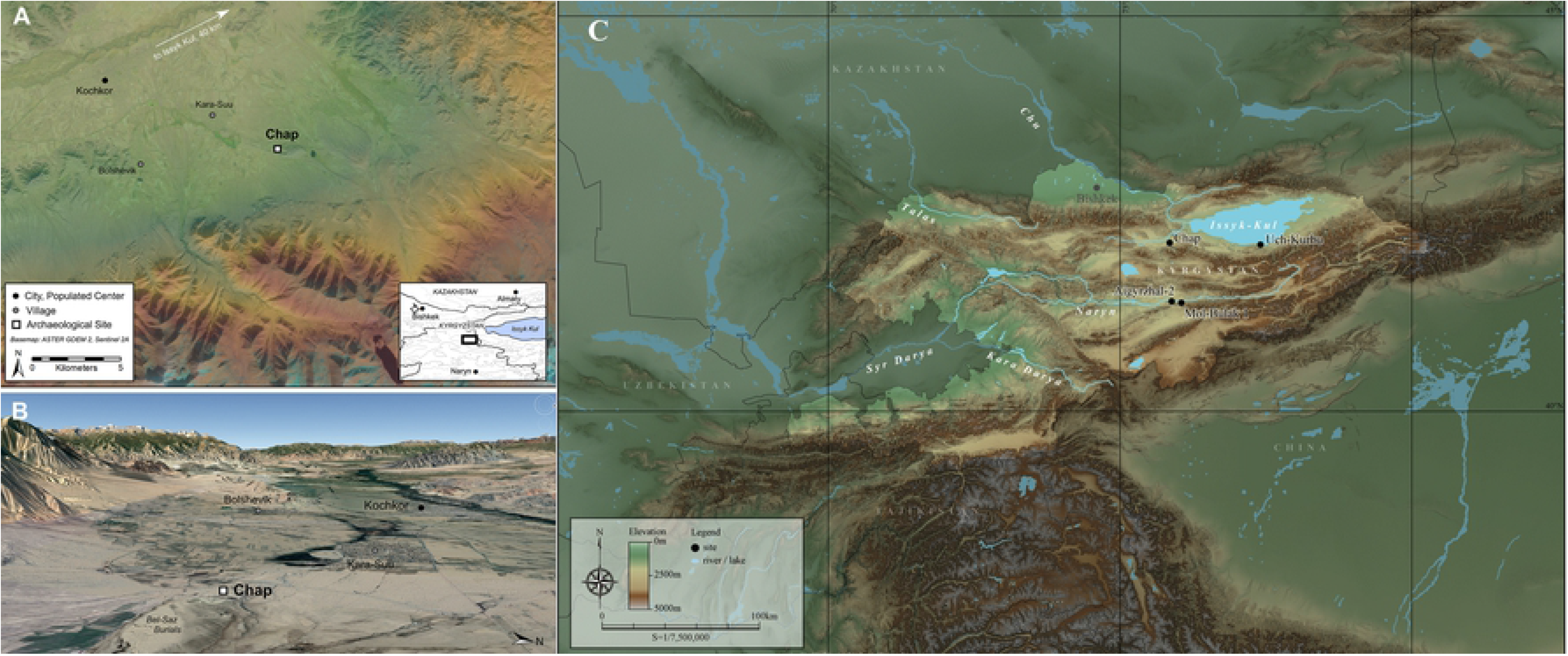
The bird’s eye view (a) and the perspective view (b) (courtesy of Google Earth) (b) of the Kochkor valley facing west, with location of the Chap site shown situated near the village of Karasu and Kochkor city. (c) The Chap site in relation to other archaeobotanically analyzed sites in Kyrgyzstan.

The Chap II site is located stratigraphically below the occupational layers of Chap I and was excavated in the summer of 2019. Chap II is separated from the final occupational strata of Chap I by a *ca* 1-m thick loess deposit. The archaeological horizon of Chap II consists of two ash pits (referred as pit 1 and pit 2), containing burned animal bones, one small fragment of pottery, and abundant carbonized plant remains. The pits are separated from one another by a sterile, 10-cm thick horizon; pit 1 is stratigraphically above pit 2, likely reflecting different formation events. The bottom portion of pit 2 reaches a depth of 260 cm above the modern surface and cradles a large, in-situ boulder. The deposition of pit 1 and 2 most likely were the result of discarded domestic waste, since the pits were not associated with discernable features that could be interpreted as being used for mortuary purposes or other commemorative practices.

## Methods

### Archaeobotanical analysis

We followed the British Heritage strategy of macrobotanical sampling by collecting at least 40 L of sediment from two different areas of each archaeological context. This sediment volume produces a more comprehensive diversity of plant taxa in archaeological deposits than that yielded from sample volumes of 10 L, which in turn, better reflects human subsistence activities [25]. Bucket flotation using a 0.3 mm screen was used to float sediment samples. In total, 136 L of sediment from the two discernible ash pits were floated: 40 L from pit 1 and 96 L from pit 2. The flotation samples were analyzed archaeobotanically at the Bioarchaeology Research Centre of Vilnius University, where they are currently archived. The stereomicroscope and the reference collection of Bioarchaeology Research Centre, including botanical seed atlases, were used to identify and quantify plant remains [26,27]. No permits were required for the described study, which complied with all relevant regulations.

### Radiocarbon dating

Radiocarbon dating of carbonized grains by accelerator mass spectrometer (AMS) ^14^C dating was performed at the 14CHRONO Centre for Climate, the Environment, and Chronology, Queen’s University Belfast. Three cereal caryopses collected from flotation samples were selected for direct dating: two wheat grains from pit 1 and one barley grain from pit 2. Radiocarbon determinations were calibrated using IntCal13 calibration curve [28], which were modelled in OxCal v. 4.3 [29] according to their stratigraphic positions relative to previously identified occupational layers yielding ^14^C dates of domesticated grains recovered from Chap I previously reported by Motuzaite Matuzeviciute et al. [24].

## Results

### Cultivated plants

The archaeobotanical assemblages of pits 1 and 2 are dominated by grains and chaff of free-threshing and glume wheats and also barleys of naked and hulled varieties. In total, 661 free-threshing wheat caryopses (*Triticum aestivum/durum*), including whole and slightly fragmented grains, were recovered. The free-threshing wheat grains have highly compact morphotypes with strikingly large variation in grain size, ranging from 6 to 2.4 mm in length, 4.2 to 1.8 mm in breadth, and 3.4 to 1.2 mm in depth. The majority of wheat grains probably belong to two free-threshing species of *T. durum* and *T. aestivum* (Fig 2). The grains of those species were counted together, yet the presence of rachis internodes belonging to bread wheat and durum wheat shows that the assemblage contains both species (Table 1) (Fig 3a, b).

**Table 1.**
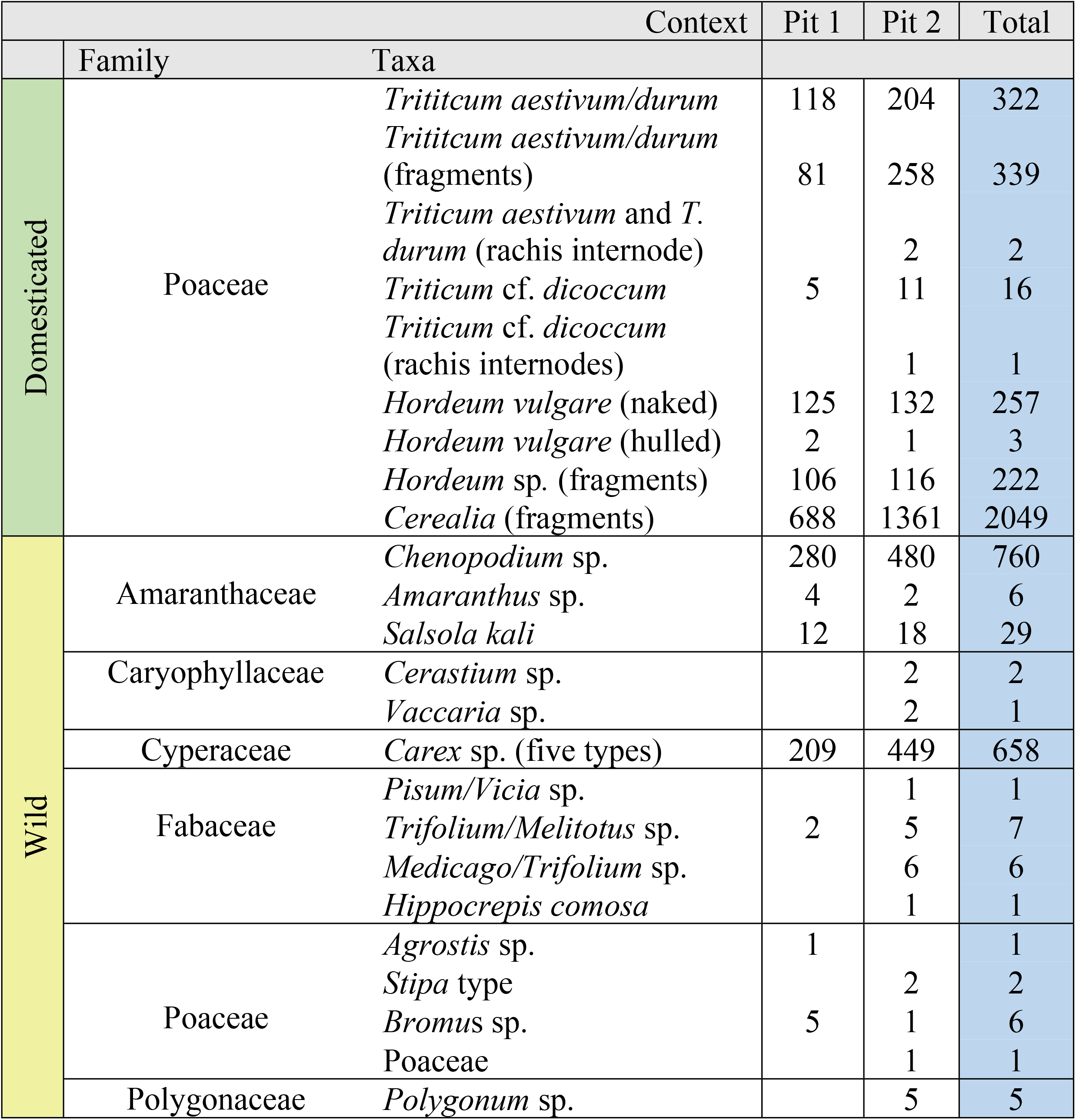
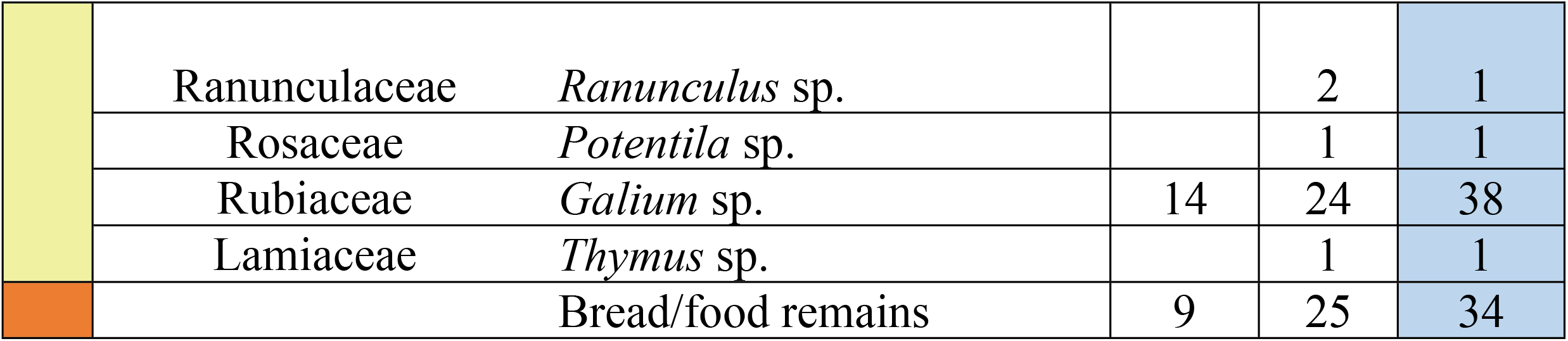
Archaeobotanical identifications from Chap II.

| Context |  |  | Pit 1 | Pit 2 | Total |
| --- | --- | --- | --- | --- | --- |
|  | Family | Taxa |  |  |  |
| Domesticated | Poaceae | *Triticum aestivum/durum* | 118 | 204 | 322 |
|  |  | *Triticum aestivum/durum* (fragments) | 81 | 258 | 339 |
|  |  | *Triticum aestivum* and *T. durum* (rachis internode) |  | 2 | 2 |
|  |  | *Triticum* cf. *dicoccum* | 5 | 11 | 16 |
|  |  | *Triticum* cf. *dicoccum* (rachis internodes) |  | 1 | 1 |
|  |  | *Hordeum vulgare* (naked) | 125 | 132 | 257 |
|  |  | *Hordeum vulgare* (hulled) | 2 | 1 | 3 |
|  |  | *Hordeum* sp. (fragments) | 106 | 116 | 222 |
|  |  | *Cerealia* (fragments) | 688 | 1361 | 2049 |
| Wild | Amaranthaceae | *Chenopodium* sp. | 280 | 480 | 760 |
|  |  | *Amaranthus* sp. | 4 | 2 | 6 |
|  |  | *Salsola kali* | 12 | 18 | 29 |
|  | Caryophyllaceae | *Cerastium* sp. |  | 2 | 2 |
|  |  | *Vaccaria* sp. |  | 2 | 1 |
|  | Cyperaceae | *Carex* sp. (five types) | 209 | 449 | 658 |
|  | Fabaceae | *Pisum/Vicia* sp. |  | 1 | 1 |
|  |  | *Trifolium/Melilotus* sp. | 2 | 5 | 7 |
|  |  | *Medicago/Trifolium* sp. |  | 6 | 6 |
|  |  | *Hippocrepis comosa* |  | 1 | 1 |
|  | Poaceae | *Agrostis* sp. | 1 |  | 1 |
|  |  | *Stipa* type |  | 2 | 2 |
|  |  | *Bromus* sp. | 5 | 1 | 6 |
|  |  | Poaceae |  | 1 | 1 |
|  | Polygonaceae | *Polygonum* sp. |  | 5 | 5 |
|  | Ranunculaceae | *Ranunculus* sp. |  | 2 | 1 |
|  | Rosaceae | *Potentilla* sp. |  | 1 | 1 |
|  | Rubiaceae | *Galium* sp. | 14 | 24 | 38 |
|  | Lamiaceae | *Thymus* sp. |  | 1 | 1 |
|  |  | Bread/food remains | 9 | 25 | 34 |

**Fig 2.**
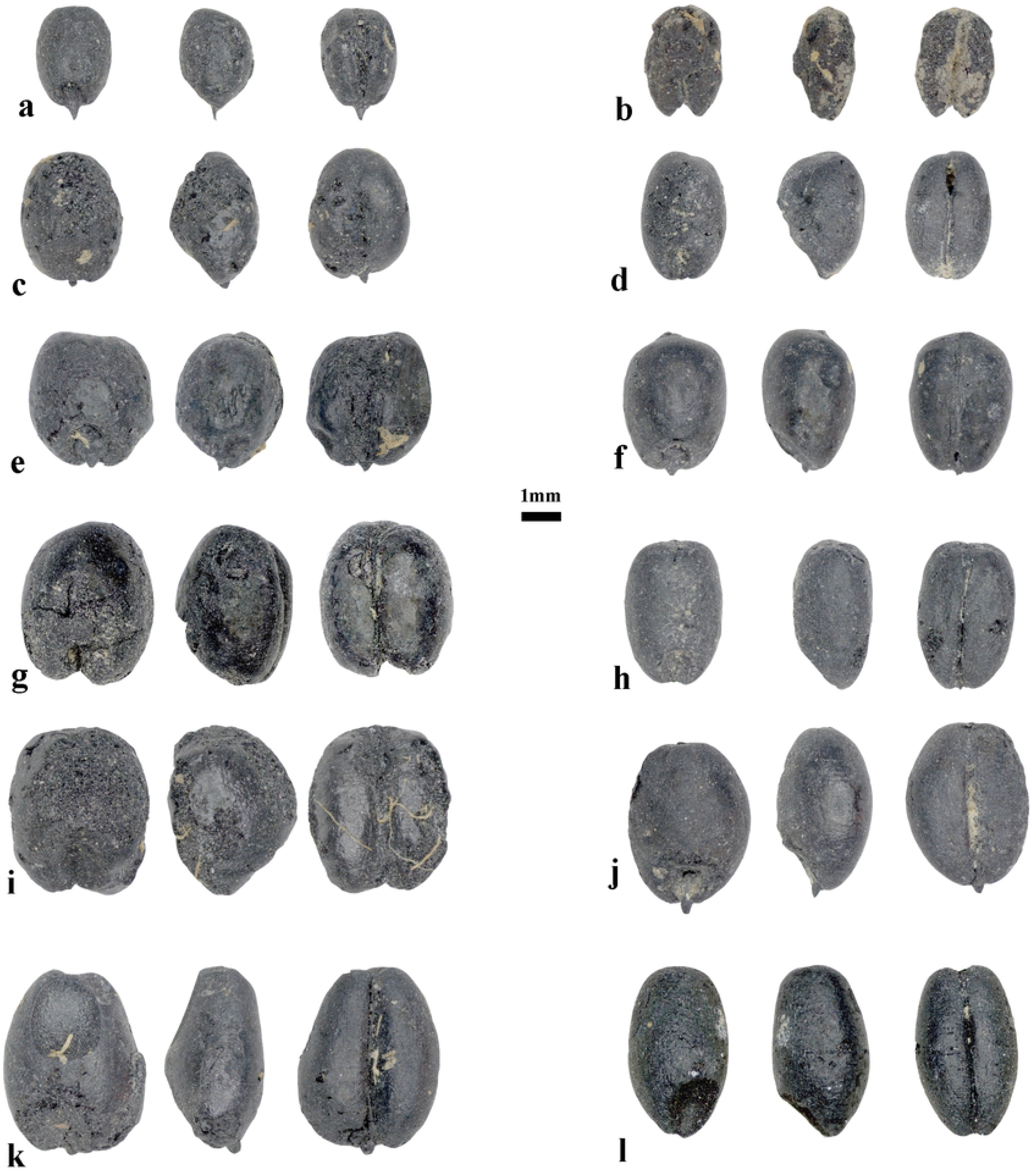
The compact wheat varieties from the Chap II site. The images of wheat show large size variation.

**Fig 3.**
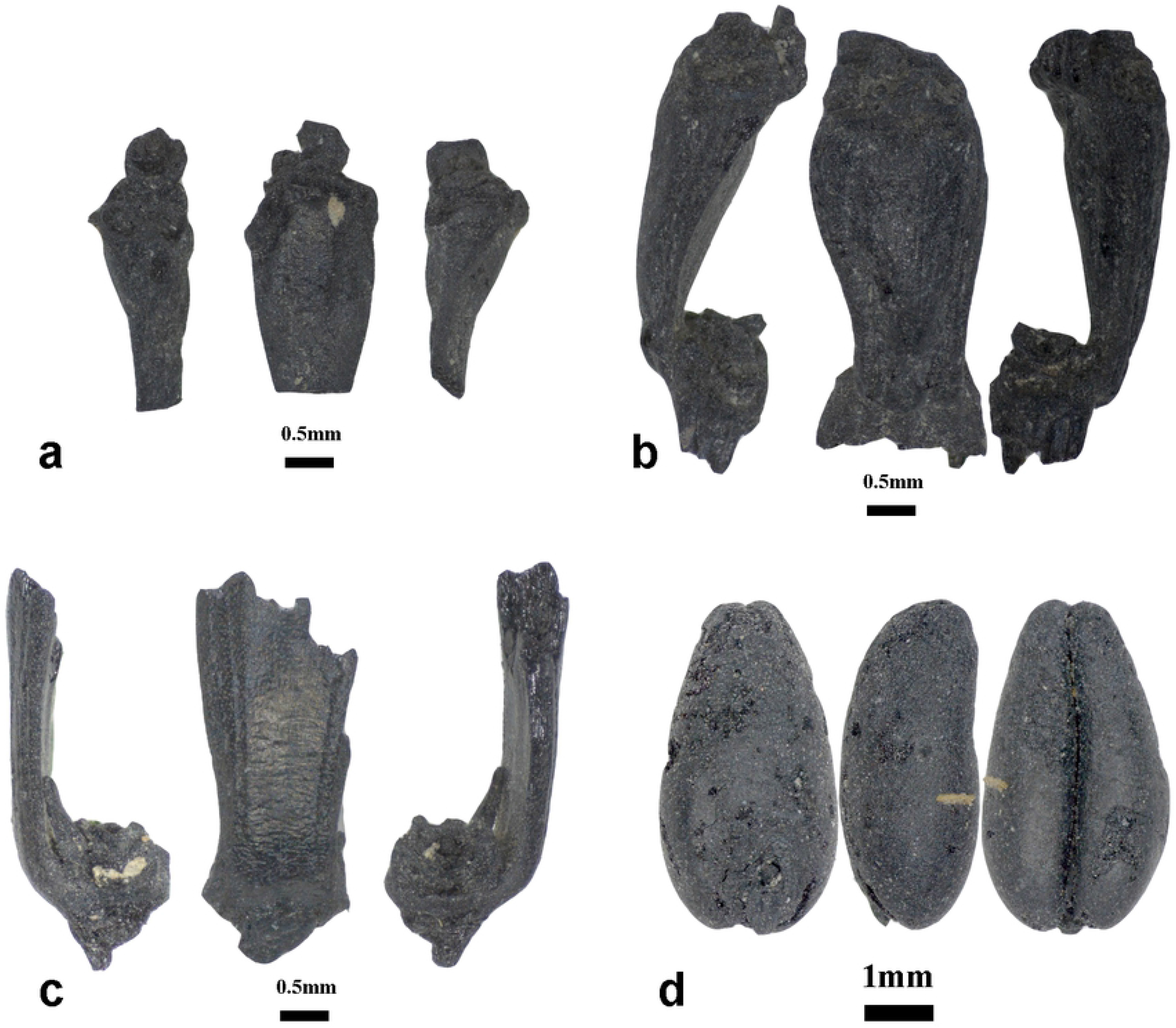
Rachis internodes of free-threshing hexaploid wheat (*Triticum* cf. *durum*) (a), *Triticum* cf*. aestivum* (b); Glume base and grains of glume wheat (*Triticum* cf. *dicoccum*) (c, d).

Sixteen grains of glume wheat, probably representing emmer wheat (*T. dicoccum*), were distinct from the free-threshing wheat grains, based on the grains having narrower widths as compared to free-threshing wheat and also exhibiting shallow, elongated and blunted apexes and embryo notches, higher dorsal ridges (keel) and slight deflections on the ventral side (Fig 3d). The recovery of one fragment of a glume base clearly shows the presence of glume wheat in the archaeobotanical assemblage of Chap II (Fig 3c).

The majority of recovered barleys belong to highly compacted varieties of *Hordeum vulgare* var. *nudum*., totaling 257 grains. As observed in wheat, the naked barley grains also vary widely in size and shape, ranging from 6.4 to 2.7 mm in length, 4.3 to 1.8 mm in breadth, and 3.5 to 1.2 mm in depth (Fig 4). Surprisingly, the measurements of barley are similar to the measurements of wheat. In fact, some of the barley grains were so strongly compacted, that differentiating them from wheat was only possible from the lateral view (Fig 4). Three barley grains were identified to hulled barley varieties. At Chap II, most barley remains are represented by symmetrical grains, probably belonging to two-or four-row varieties. Counting partial and whole grains together, both pits yielded 482 grains of barley. Due to fragmentation, 2040 grain remains were identified as “Cerealia” type.

**Fig 4.**
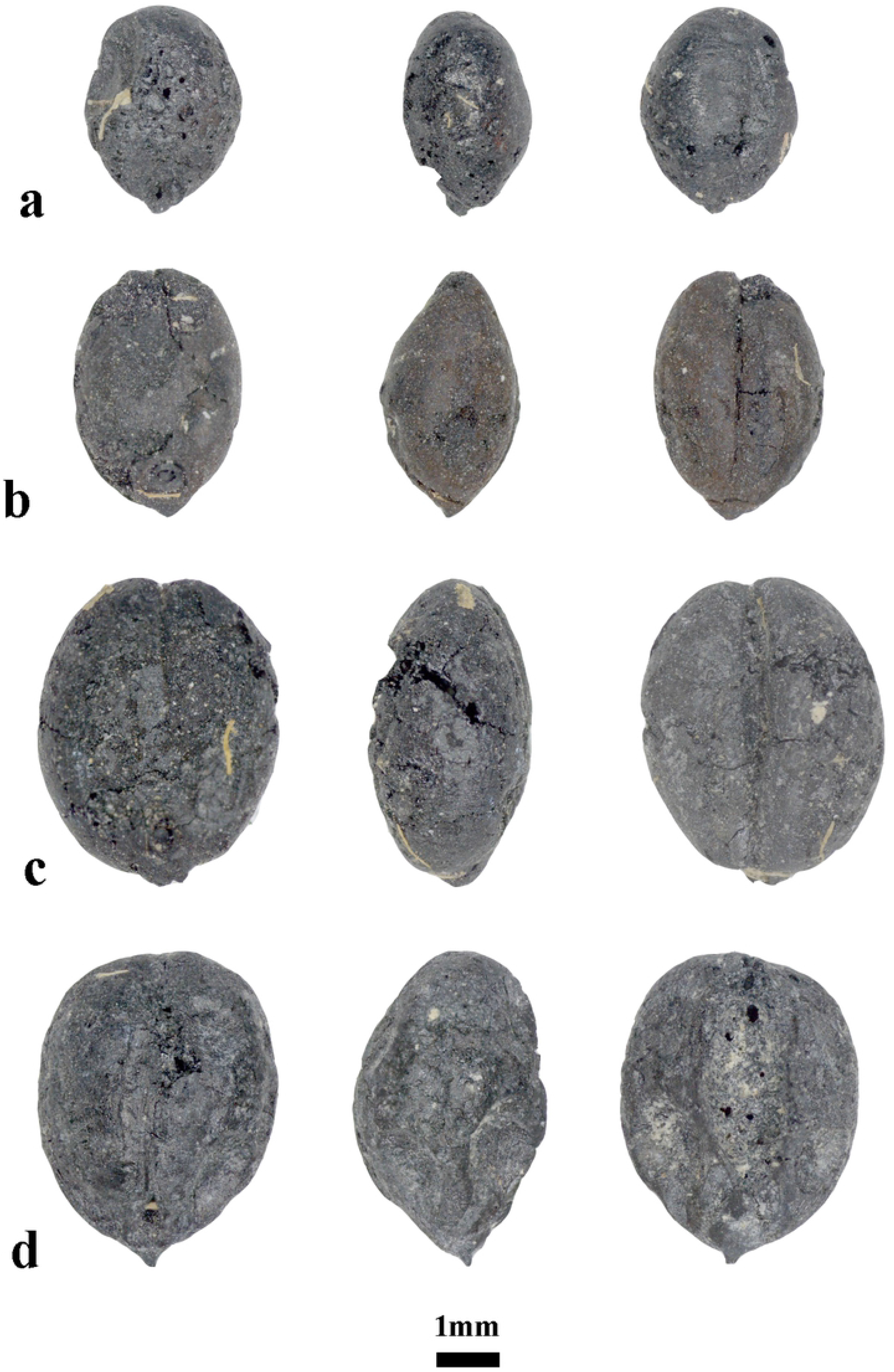
Highly compact morphotypes of naked barley from Chap II.

It was surprising that no remains of East Asian crops were recovered from the flotation samples, such as broomcorn or foxtails millet, which were present at Chap I [24]. Interestingly, 34 carbonized food fragments, up to 9.5 mm in size, were also identified. These remains were characterized by porous or amorphous-like matrices and a burned, lumpy conglomerate, some containing cereal bran fragments. These charred food pieces probably represent the remains of bread or porridge. The food fragments were not analyzed in this study but will be the subject of future research.

### Wild plants

The most common seed remains in the wild plant assemblage were from *Chenopodium* sp. (goosefoot), represented by 760 carbonized grains (Table 1). *Chenopodium* plants normally represent ruder weeds that grow in nitrogen-rich, former domestic spaces [30,31]. The second most abundant wild seeds were from *Carex* sp. (sedges), totaling 658 carbonized grains (Table 1). Five morphotypes of *Carex* caryopses were identified, which likely all belong to different species (Fig 5). Most *Carex* plants grow in wetland ecosystems and tolerate saline waters [32]. The presence of sedge seeds among the domesticated grains at Chap II could mean that crops were grown in close proximity to sedges, possibly along irrigated channels or mountain streams. The other wild species are represented by smaller seed counts. The majority of them reflect open meadow, arable and possibly irrigated landscapes, such as *Salsola kali*, *Polygonum* sp. or *Galium* sp. (Table 1). Moreover, the majority of taxa are not found in the undergrowth of forests [33,34], which helps illustrate a local environment surrounding Chap II that was devoid of trees.

**Fig 5.**
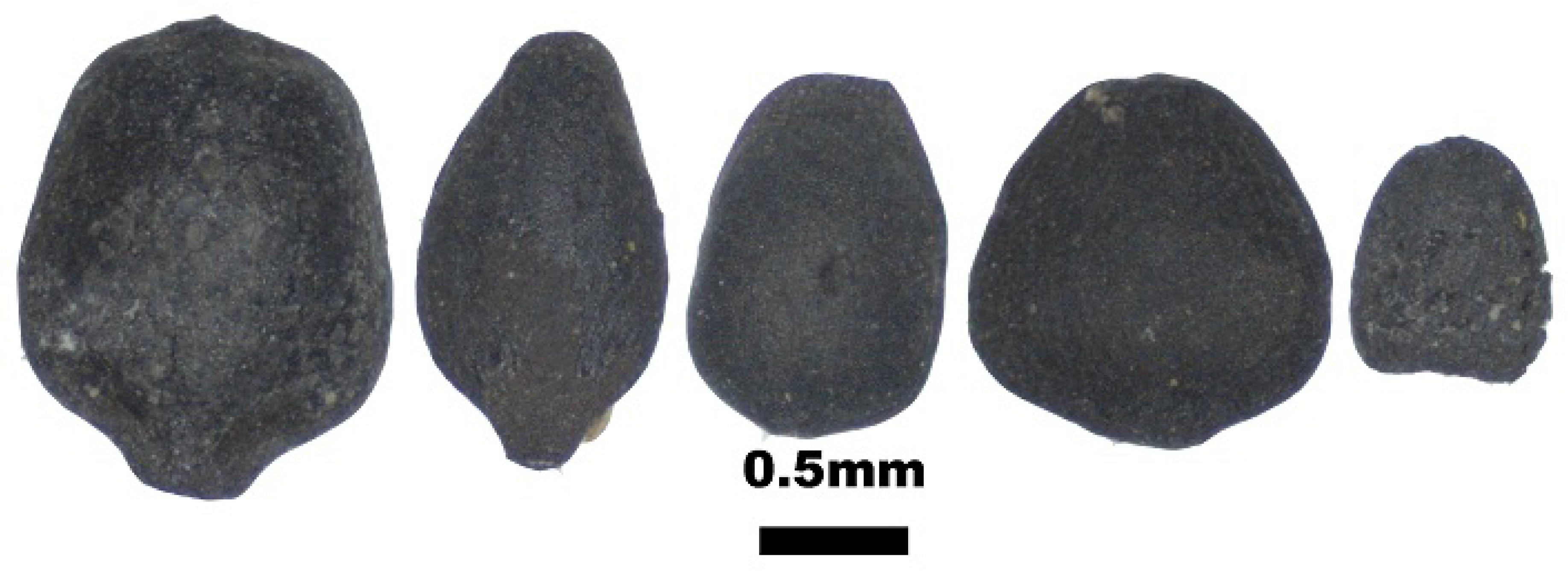
The five types of *Carex* sp. (sedges) found at Chap II.

### Radiocarbon dating

Three radiocarbon determinations were measured on two individual wheat grains from pit 1 and one individual barley grain from pit 2. Showing consistency with the relative stratigraphic position of pit 2 below pit 1, the modelled date of the barley grain from pit 2 is 2467-2292 cal BCE, while the wheat grains from pit 1 together provide a modelled date of 2402-2047 cal BCE (Table 2; Fig 6). Overall, the two pits date to nearly 1500 years before the occupation of Chap I, raising the possibility that the site was abandoned for a considerable amount of time during the second millennium BCE (Fig 6).

**Table 2.**
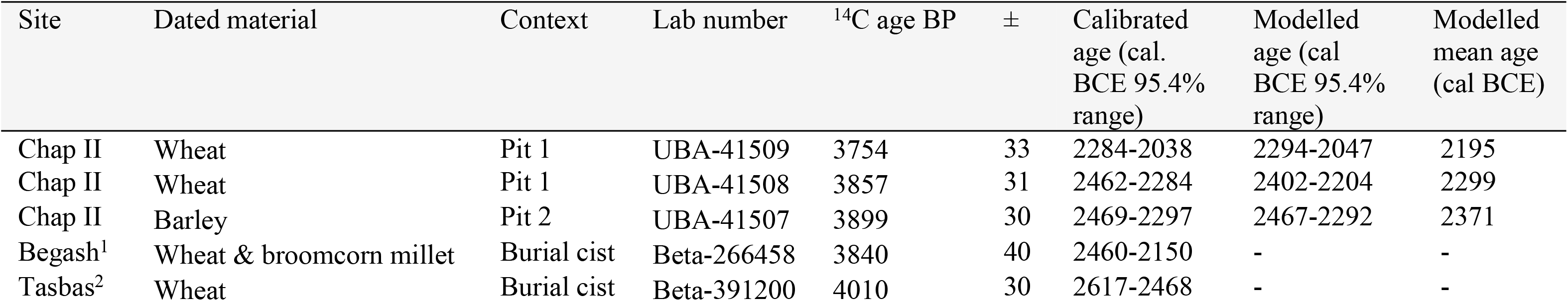
AMS radiocarbon dates measured on macrobotanical remains recovered from Chap II. Previously published radiocarbon dates taken on domesticated grains from Begash [7] and Tasbas [23] are also shown to help contextualize the chronology of Chap II.

| Site | Dated material | Context | Lab number | <sup>14</sup>C age BP | ± | Calibrated age (cal. BCE 95.4% range) | Modelled age (cal BCE 95.4% range) | Modelled mean age (cal BCE) |
| --- | --- | --- | --- | --- | --- | --- | --- | --- |
| Chap II | Wheat | Pit 1 | UBA-41509 | 3754 | 33 | 2284-2038 | 2294-2047 | 2195 |
| Chap II | Wheat | Pit 1 | UBA-41508 | 3857 | 31 | 2462-2284 | 2402-2204 | 2299 |
| Chap II | Barley | Pit 2 | UBA-41507 | 3899 | 30 | 2469-2297 | 2467-2292 | 2371 |
| Begash<sup>1</sup> | Wheat & broomcorn millet | Burial cist | Beta-266458 | 3840 | 40 | 2460-2150 | - | - |
| Tasbas<sup>2</sup> | Wheat | Burial cist | Beta-391200 | 4010 | 30 | 2617-2468 | - | - |

**Fig 6.**
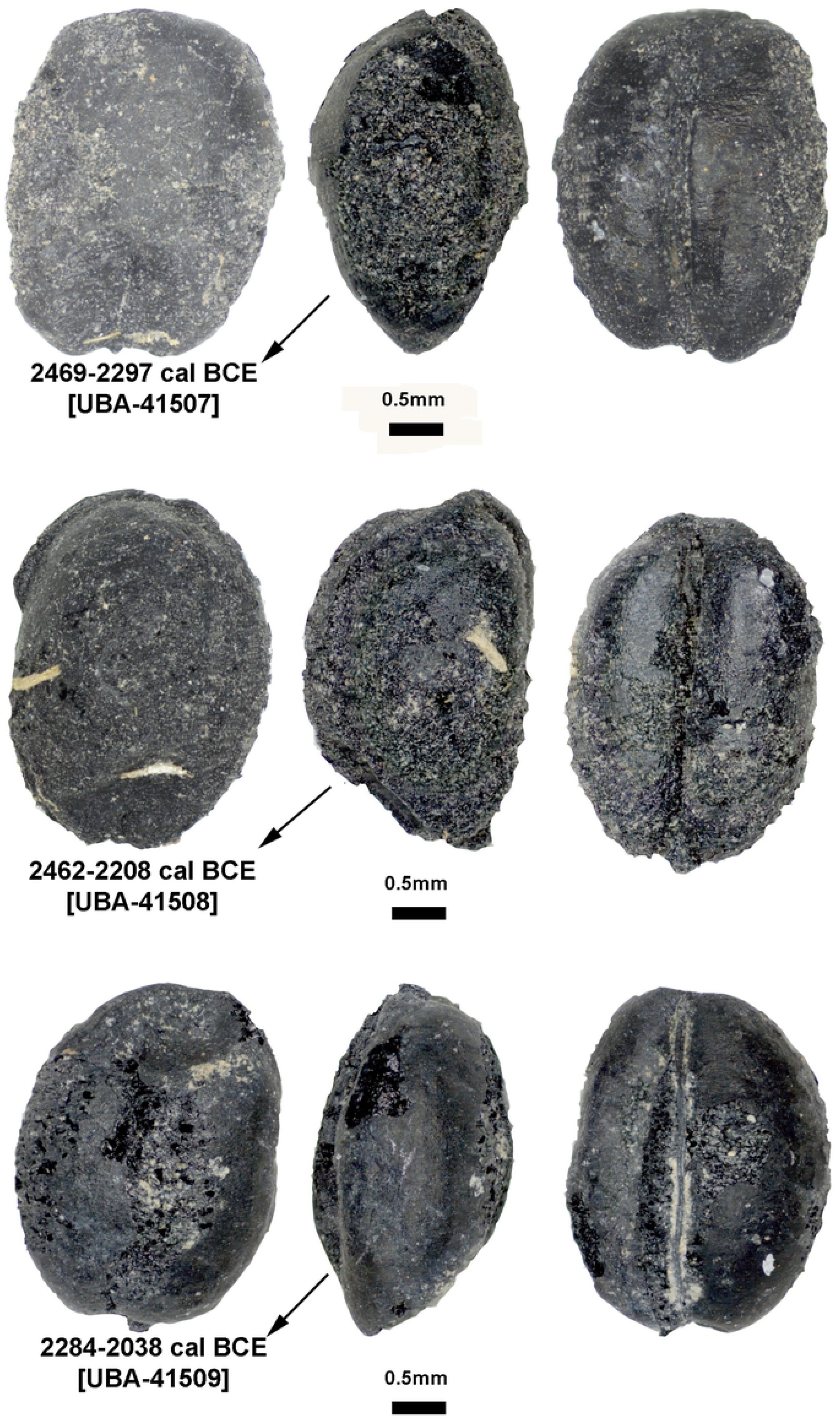
The images of directly radiocarbon dated naked barley from pit 2 (top) and free-threshing wheat grains (middle and bottom) from pit 1 with listed calibrated dates from Chap II site.

**Fig 7.**
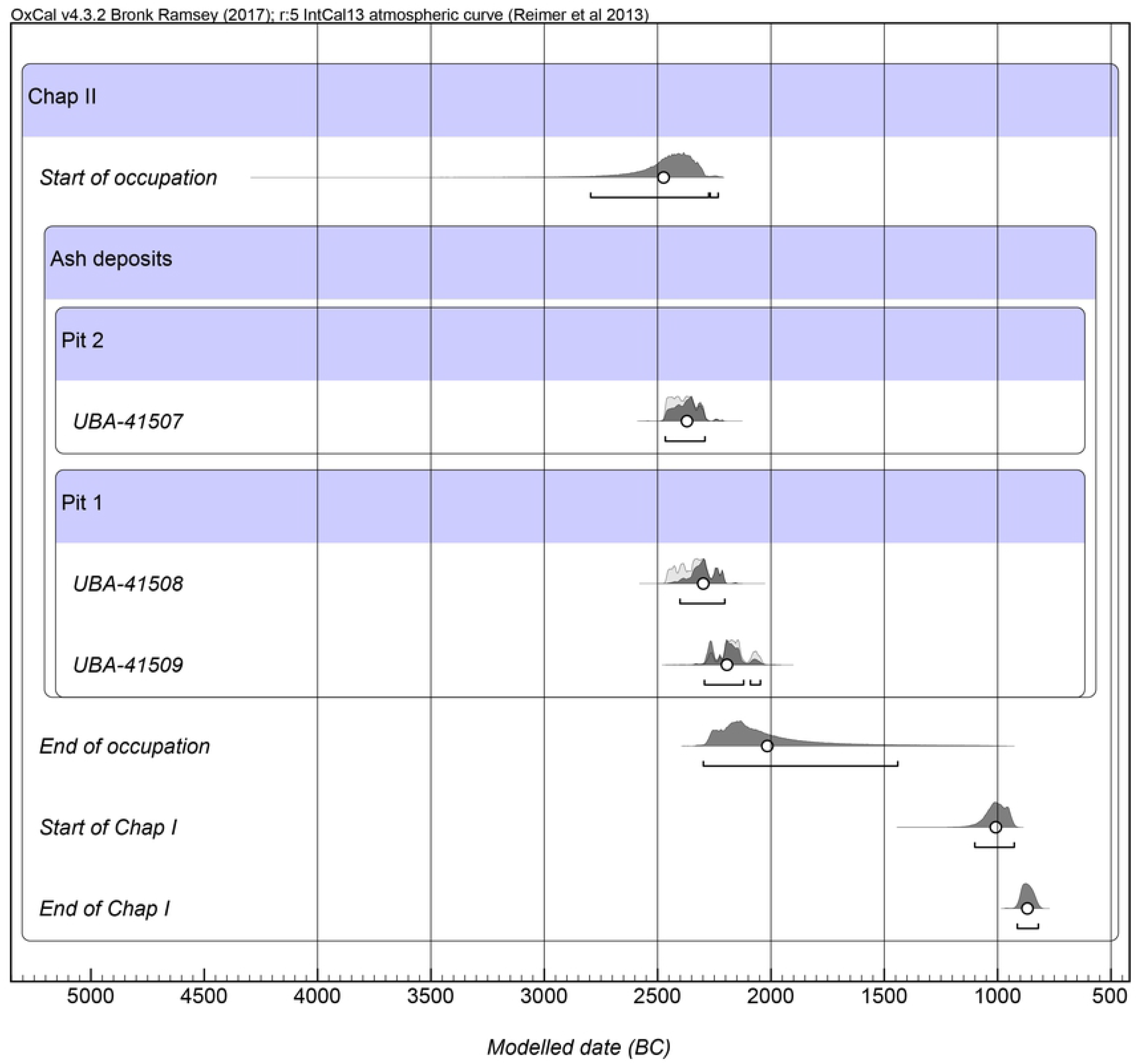
Modelled radiocarbon dates measured on wheat barley grains from Chap II in relation to the occupational period of Chap I previously reported by Motuzaite Matuzeviciute et al. [24].

## Discussion

Dated to the middle to second half of the 3^rd^ millennium cal BCE, the archaeobotanical assemblage of Chap II represents the earliest and the most abundant assemblage of cereal and wild plant macrobotanical remains in eastern Central Asia and is further notable for its recovery at 2000 m.a.s.l. Contrary to previous suggestions that an initial eastward dispersals of wheat and barley may have diverged from one another upon reaching the Inner Asian Mountain Corridor [4], the early co-occurrence of both wheat and barley in similar proportions at Chap II suggests that these crops spread into the highlands of Central Asia as a package. This scenario indicates that people engaging in plant cultivation in the Tien Shan were equally invested in varieties of wheat and barley that already had been adapted to high elevations, where short growing seasons and low temperatures impose considerable difficulties for farmers. The biodiversity of crops, reflected in both the compact grain morphotypes, large variation in cereal grain sizes, and relative abundance of taxa, present at Chap II illustrates subsistence strategies that likely favored agricultural resilience by safeguarding crop production from extreme and uncertain weather conditions characteristic of montane environments. However, it is notable that the archaeobotanical assemblage of Chap II lacks Chinese domesticates of millet, which provide agriculturalists with additional flexibility in production on account of millets having short growing seasons and drought tolerance. The fact that broomcorn millet was recovered at Begash located at 900 m.a.s.l. [7] and likely cultivated there and at Dali located at 1500 m.a.s.l. [16], both in the neighboring region of Semirech’ye (southeastern Kazakhstan), had previously implied that millet may have also reached the central Tien Shan by the end of the third millennium BCE. The definitive absence of millet at Chap II may indicate that 1) interaction networks reaching into the northern stretches of the Inner Asian Mountain Corridor did not connect people living in the central Tien Shan or 2) an elevational ceiling for successful cultivation below 2000 m.a.s.l. existed for millet cultigens in the third millennium BCE. During the occupation of Chap I 1500 years later, millets formed an important component of the local crop repertoire, potentially overcoming prior environmental constraints [35].

The archaeobotanical assemblage at Chap II site is dominated by domesticates of southwest Asian origin: free-threshing wheats (possibly *T. durum* and *T. aestivum*) and both naked and hulled barleys. The free-threshing wheat grains correspond to compact wheat forms noted in the Dzhungar Mountains of Semirech’ye [6,18,19]. Furthermore, archaeobotanical research in Kyrgyzstan identified compact wheat forms at the sites of Uch Kurbu, Aigyrzhal-2, and Mol Bulak located in the highland regions (Fig 1c) [15,17], which suggests that compact wheat morphotypes are adapted to high elevations. Other researchers have also noted the suitability of compact wheats for agriculturalists in marginal environments of south and east Asia, such as monsoonal areas of India and the highlands of the Tibetan plateau [14,36]. It has been argued that, compared to non-compact forms, compact varieties of wheat exhibit greater standing power against extreme weather conditions, such as strong winds and intense rains that can cause lodging and stem breakage [15]. In addition to compact forms, large variation in wheat grain size was also noted for the wheats at Chap II, ranging from 6.0 to 2.4 mm in length, 4.2 to 1.8 mm in breadth and 3.4 to 1.2 mm in depth. This wide variation in grain size may be the result of environmental factors that crop plants were exposed to during grain development. Previous experimental research has shown that plants experiencing physiological stress from differences in water availability, ambient temperature, or amount of nutrients during grain filling resulted in grain size diminution, due to interference in the deposition of carbohydrates in the grains [37–42]. In addition, Reed [43] also showed that grains from a single ear of spelt wheat exhibited a wide variety of sizes, indicating that variability in grain size can also be due to heterogeneous ripening times of grains on the ear.

It is noteworthy to point out that the biometrical data for wheat grains recovered from Bronze and Iron Age sites in Kyrgyzstan substantially overlap with that of wheat grains from second millennium BCE sites in north western China [14,15]. In eastern regions of monsoonal China, however, ancient wheat grains show dramatically smaller sizes and follow a more uniform size distribution, which suggest that the eastward spread of wheat was associated with an overall convergence to diminutive grains [14]. Further research is needed in order to understand whether the reduction in wheat grain size in monsoonal China was influenced by climate, human choice, or the geographical origins of particular wheat morphotypes.

The presence of two types of free-threshing wheat rachis internodes (Fig 3) and the large diversity of wheat grain morphotypes found at Chap II and at other sites in Kyrgyzstan, such as Argyrzhal-2 and Uch-Kurbu [17], suggest that probably both tetraploid and hexaploid free-threshing wheat were cultivated. In China, archaeobotanical research mainly reports grain morphology, thus offering opportunities for future analysis of rachis internodes to test whether both tetraploid and hexaploid free-threshing wheats were cultivated there and, possibly, to resolve the timing and dispersal routes of these species.

The discovery of glume wheat at Chap II site during early phases of southwest Asian crop dispersals to the central Tien Shan is also important for understanding the limits of its eastward spread. Current evidence suggests that the cultivation of glume wheat did not reach the present-day territory of China at the initial stage of southwestern crop dispersal during the third-second millennia BCE. The free-threshing wheats were selected instead of glume wheats as they dispersed eastwards from centers of domestication in southwest Asia [44]. Most ancient Chinese wheat remains are hexaploid free-threshing *Triticum aestivum* [45–47], in contrast to the types of wheat recovered from Neolithic and Bronze Age sites in southwestern Asia, where agriculture was mainly based on glume wheat [44].

A review by Stevens et al. [44] reported modern glume wheat varieties (*T. aestivum* var. *tibetianum* JZ Shao) in western China with limited distribution in Yunnan, Tibet, and Xinjiang, in addition to a glume wheat variety in Yunnan (*Triticum aestivum* subsp. *yunnanense* King ex SL Chen). Notably, the rachis internode of the glume wheat recovered from Chap II does not resemble that of typical glume wheat belonging to *T. dicoccum* found in Europe. Comparing the glume wheat internode bases from Chap II with those previously reported at the *Linearbandkeramik* site of Ratniv-2 site located in western Ukraine [48], the curve between glume and glume base at Chap II is straighter and *ca* 25% larger. Therefore, glume wheat at Chap II could represent local Central Asian variety or be related to modern Chinese varieties mentioned above. A few grains of glume wheats (but no chaff) have also been recently reported from Kanispur site in Kashmir, dating by associated charcoals to a similar period as Chap II in the third millennium BCE [49]. Further research is needed to understand why glume wheats were used but later disappeared from the package of cultivars among mountain communities of eastern Central Asia.

The archaeobotanical assemblage of Chap II is dominated by naked varieties of barley, which became commonly cultivated across the Tibetan plateau [36,50]. The reason for the dominance of naked forms of barley in the highlands of Asia is not clear, but it has been suggested that cultural preference was the main factor for selecting naked over hulled barleys [50]. However, naked barley is better adopted to highland environments, especially those characterized by strong continental climates [51,52]. In mountain cultivation areas, naked varieties produce higher grain yields, generate more overall biomass, and mature faster than hulled varieties [53].

The finding of highly compacted barley at Chap II is also particularly interesting as it reveals what morphotypes were selected by early farmers inhabiting mountainous landscapes and gives insight on crop adaptation in these highland regions during the third millennium BCE. Future studies comparing metrical variation of lowland and highland barley grains and chaff could lead to a better understanding of how barley varieties were advantageous in certain environmental niches. Knüpffer [52] note that landraces of modern naked barleys collected from the high-mountain regions of Central Asia are short with thick, lodging straw and have grains that are more compact and spherical than those of taller barley varieties.

It has been suggested that plump forms of wheat and six-row naked barley, in particular, are water-demanding crops that likely required irrigation or high amounts of rainfall for successful cultivation in western Central Asia and the Near East [54]. The possibility that the inhabitants of Chap II used irrigation in the Kochkor valley cannot also be ruled out. It is plausible that irrigation technologies spread together with the first farming communities to the central Tien Shan and further eastwards to China [55]. The occupation of Chap II during the second half of the third millennium BCE coincides with the Subboreal climate period of dry and cold conditions [56,57]. People may have been motivated to seek out high-mountain water sources, such as perennial springs, when the lowlands were characterized by water shortages due to glacial melt waters decreasing as precipitation diminished and ice sheets expanded. Moreover, herding domesticated animals would have also motivated people to seek highland settlements near rich, montane pastures, where they could have integrated pastoralist and plant cultivation strategies through foddering with agricultural byproducts and dung manuring [16]. One of the dominant species of wild plants at Chap II is *Carex* sp. (sedges), that normally grow in wetland environments, which suggests widespread availability of water-saturated soils. Although the archaeological deposition of sedges at Chap II remains may be the result of animal foddering or craft production, the finding of sedges among cultivated crops at the Bronze Age Kültepe-Kanesh archaeological site in Turkey led researchers there to suggest that crops were watered by irrigation channels [58]. Currently, sedges grow along modern irrigation channels in the Kochkor valley. At Chap II, the fields were probably located in close proximity to water sources, either irrigated channels or natural streams that could be easily redirected. Stable isotopic analysis of the grains recovered at Chap II is needed to examine watering regimes for local plant cultivation, as has been previously applied elsewhere [59,60].

Wild plants and ruderal weeds recovered from Chap II are dominated by *Chenopodium* sp., which together with *Galium* sp., normally inhabit nitrogen-rich domestic settings modified by intensive pastoralist herding and corralling [30]. The few faunal skeletal remains were recovered from Chap II (~25 specimens), likely represent domesticated ruminant species. The presence of domesticated sheep and goats dated to ~2700 cal BCE were identified by mitochondrial DNA at Dali located in the Dzhungar Mountains of southeastern Kazakhstan [16]. It is possible that pastoralist subsistence spread northward through the Tien Shan mountains along the Inner Asian Mountain Corridor earlier than the occupation of Chap II [61], although additional excavation of a wider area of Chap II will aid in the recovery of additional faunal specimens. Judging from the rest of the wild plant and weed taxa, it is difficult to be certain whether the recovered crops from Chap II were sown in the autumn or spring. Most of the abundant wild plants, such as *Chenopodium* sp. and *Galium* sp., are attributed to summer annual plants, which hints to spring-sown varieties of wheat and barley. Nonetheless, the subsistence strategies in effect at Chap II during the second half of the third millennium BCE likely involved intensive investment in plant cultivation of numerous cultigens with varied cultivation schedules, in addition to people also engaging in pastoralist herding involving varied mobility strategies.

## Conclusions

The Chap II site yielded the largest crop assemblage dated to the third millennium BCE between Pamir, Tien Shan, and Altai mountains, comprising thousands of cultivated cereal remains with the presence of two species of free-threshing wheat (bread wheat and durum wheat), glume wheat, and hulled and naked barley. Radiocarbon determinations derived directly from wheat and barley grains show that Chap II was occupied between 2467-2047 cal BCE. Analysis of the accompanying weed taxa recovered from Chap II hints at an open landscape where pastoralist herding likely took place. The dominance of wetland plants in the assemblage also suggests that cultivated crops may have been irrigated, which would imply that this technology spread hand in hand the cultivation of southwest Asian crops. Finally, the chaff and grain remains of glume wheat represents the first time this species was recovered in the most eastern regions of Central Asia, which critically invigorates a new discussion about why and where glume wheats became filtered out of crop repertories as other southwest Asian crops spread to the eastern parts of Eurasia.

## Acknowledgements

This research is funded by the European Social Fund according to the activity ‘Improvement of researchers’ qualification by implementing world-class R&D projects’ of Measure No. 09.3.3-LMT-K-712. We would like to acknowledge 14Chrono Centre at Queen’s University Belfast for providing radiocarbon dates at the collaborator’s rate.

